# REDD1 is a determinant of the sensitivity of renal cell carcinoma cells to autophagy inhibition that can be therapeutically exploited by targeting PIM kinase activity

**DOI:** 10.1101/2024.10.29.620949

**Authors:** Jennifer S. Carew, Claudia M. Espitia, Sruthi Sureshkumar, Maria Janina Carrera Espinoza, Madison E. Gamble, Wei Wang, Benjamin R. Lee, Steffan T. Nawrocki

## Abstract

**Purpose:** Repurposing FDA approved drugs with off-target autophagy inhibition such as chloroquine/hydroxychloroquine (CQ, HCQ) has produced modest anticancer activity in clinical trials, due in part, to a failure to define predictive biomarkers that enable the selection of patients that best respond to this treatment strategy. We identified a new role for REDD1 as a determinant of sensitivity to autophagy inhibition in renal cell carcinoma (RCC).

**Experimental Design:** RNA sequencing, qRT-PCR, immunoblotting, gene silencing, knockout and overexpression studies revealed that REDD1 expression is a key regulator of cell death stimulated by autophagy inhibitors. Comprehensive *in vitro* and *in vivo* studies were conducted to evaluate the selectivity, tolerability, and efficacy of the PIM kinase inhibitor TP-3654 and CQ in preclinical models of renal cell carcinoma (RCC). Markers of autophagy inhibition and cell death were evaluated in tumor specimens.

**Results:** Transcriptomic analyses identified REDD1 (*DDIT4*) as a highly induced gene in RCC cells treated with the PIM kinase inhibitor TP-3654. Focused studies confirmed that PIM1 inhibition was sufficient to induce REDD1 and stimulate autophagy through the AMPK cascade. *DDIT4* knockout and overexpression studies established its mechanistic role as a regulator of sensitivity to autophagy inhibition. Inhibition of autophagy with CQ synergistically enhanced the i*n vitro* and *in vivo* anticancer activity of TP-3654.

**Conclusions:** Our findings identify REDD1 as a novel determinant of the sensitivity of RCC cells to autophagy inhibition and support further investigation of PIM kinase inhibition as a precision strategy to drive sensitivity to autophagy-targeted therapies through REDD1 upregulation.

**Translational Relevance:** PIM kinases are overexpressed in renal cell carcinoma (RCC) and other malignancies. Here we show that targeting PIM1 significantly upregulates REDD1/*DDIT4* expression, resulting in inhibition of mTOR and autophagy activation. REDD1 induction was determined to be a major factor that regulates the sensitivity of RCC cells to autophagy inhibition. Comprehensive *in vitro* and *in vivo* studies in preclinical models of RCC demonstrated that the PIM kinase inhibitor TP-3654 induced REDD1-mediated autophagy and synergistically sensitized cells to autophagy inhibition. Our findings define a new role for REDD1 as a determinant of the sensitivity of RCC cells to autophagy inhibition and demonstrate that antagonizing PIM kinase activity is a precision approach to augment REDD1 levels and potentiate the therapeutic benefit of targeting the autophagy pathway.

## Introduction

Renal cell carcinoma (RCC) accounts for 85% of kidney cancers with clear cell RCC (ccRCC) being the most common subtype representing approximately 75% of all cases (1). Loss of expression of the von Hippel-Lindau (VHL) tumor suppressor occurs in 70% of sporadic ccRCC patients and causes stabilization of hypoxia-inducible factors (HIFs) (2). HIFs alter the cellular environment by activating target genes involved in angiogenesis and metabolism (3). Multi-tyrosine kinase inhibitors including sunitinib and cabozantinib and mammalian target of rapamycin (mTOR) inhibitors such as temsirolimus and everolimus have demonstrated efficacy for the treatment of ccRCC. Immune checkpoint inhibitor therapies targeting programmed cell death protein 1 (PD-1) and cytotoxic T lymphocyte-associated protein 4 (CTLA-4) have also recently demonstrated significant clinical activity. Despite the benefit yielded by these agents, drug resistance is a major obstacle that continues to limit successful clinical outcomes especially for patients with metastatic ccRCC (4). While autophagy has been established as a key pro-survival mechanism that promotes tumor progression, metastasis, and drug resistance, specific biomarkers that predict sensitivity or define subsets of patients that may benefit most from autophagy inhibitors have not been validated (5–7). As various autophagy inhibitors enter clinical testing, a better understanding of the factors that drive sensitivity to autophagy inhibition may yield new therapeutic approaches to enhance their anticancer activity.

The PIM kinases are a family of serine/threonine kinases that have been associated with tumorigenesis and drug resistance (8–11). The kinases have an ATP binding pocket, an active site, and a kinase domain, but are constitutively active due to their lack of regulatory domains. PIM kinases regulate cell cycle progression by directly phosphorylating p21, p27, and Cdc25C and suppress apoptosis via phosphorylation of the pro-apoptotic protein Bad (11–14). Overexpression of PIM1 has been reported in hematological malignancies and in many solid tumors (9–11). In ccRCC specifically, PIM1 overexpression is correlated with disease progression and metastasis, which suggests that blocking PIM1 kinase activity may be a promising therapeutic approach for patients that do not benefit optimally from conventional treatment (15,16). Consistent with these reports, we demonstrated that PIM1 is overexpressed in ccRCC models and its inhibition yields promising anticancer activity, is well tolerated, and potentiates standard chemotherapy (17).

TP-3654 is a second-generation PIM kinase inhibitor that has entered into early phase clinical trials (18–21). Here we report on its preclinical efficacy and mechanism of action in models of ccRCC. Our investigation identified REDD1 as a highly upregulated gene induced by TP-3654 treatment. Further investigation demonstrated that the specific pharmacodynamic induction of REDD1 expression is a key determinant of the sensitivity of ccRCC cells to autophagy inhibition. Co-targeting PIM kinase and autophagy led to synergistic anticancer activity in both *in vitro* and *in vivo* models of RCC. Our collective data demonstrate that REDD1 is a novel biomarker of the sensitivity of RCC cells to autophagy inhibition. Targeting PIM1 offers a precision approach to heighten REDD1 expression and synergistically augment the efficacy of the autophagy inhibitor CQ. Clinical investigation of this combination therapeutic strategy for patients with advanced RCC is warranted.

## Materials & Methods

### Cells and cell culture

A498, 786-O, Achn, Caki-1, and Caki-2 cells were purchased from ATCC (Manassas, VA). RCC4 cells were obtained from Dr. Sunil Sudarshan (University of Alabama at Birmingham, Birmingham, Alabama). RCC cells were cultured with medium supplemented with 10% FBS at 37°C with 5% CO_2_ as previously described (5). Human normal renal proximal tubule epithelial cells (RPTEC) were purchased from ATCC and cultured in Renal Epithelial Cell Basal Medium plus one Renal Epithelial Cell Growth Kit. HAP1 parental and *DDIT4*^−/−^ cells were obtained from Horizon Discovery (Cambridge, UK). Cell lines were authenticated by the source banks using DNA profiling techniques.

### Chemicals and reagents

Reagents were obtained from the following sources: propidium iodide (PI), bafilomycin A1, 3-(4,5-dimethylthiazol-2-yl)-2,5-diphenyltetrazolium bromide (MTT), chloroquine (CQ), and anti-β-tubulin (#T7816) antibody were purchased from Sigma (St. Louis, MO). TP-3654 was obtained from SelleckChem (Houston, TX). Antibodies were purchased from: anti-cleaved PARP (#9541), anti-cleaved caspase-3 (#9661), anti-PCNA (#2586), anti-p-AMPK (Thr172) (#2535), anti-AMPK (#5831), anti-TXNIP (#14715), anti-p-mTOR (Ser2448) (#5536), anti-mTOR (#2983), anti-p-p70S6K (Thr389) (#9234), anti-p70S6K (#2708), anti-p-4E-BP1 (Thr37/46) (#2855), anti-4E-BP-1 (#9644), anti-ATG7 (#8558), anti-PIM1 (#3247), anti-p27 (#3686), anti-p-BAD (Ser112) (#5284), anti-BAD (#9239), anti-p-c-MYC (Ser62) (#13748), anti-c-MYC (#18583), and anti-LC3B (#3868) (Cell Signaling, Danvers, MA), anti-p62 (#ab91526) (Abcam, Cambridge, UK), and anti-REDD1 (#10638-1-AP) (Proteintech, Rosemont, IL). Donkey anti-rabbit and sheep anti-mouse horseradish peroxidase (HRP) were obtained from Amersham (Pittsburgh, PA). Goat anti-rabbit Alexa-Fluor 594, goat anti-rabbit Alexa-Fluor 488, and Prolong Gold antifade with DAPI was purchased from ThermoFisher (Waltham, MA). Goat anti-rabbit HRP and Rat anti-mouse IgG2a-HRP tagged secondary antibodies were obtained from Jackson ImmunoResearch Laboratories (West Grove, PA).

### RNA Sequencing

RCC cells were treated with 3 μM TP-3654 for 24 h. Total RNAs were isolated using the RNeasy Plus Mini Kit (Qiagen, Germantown, MD) and treated with TURBO DNA-*free*™ Kit (Applied Biosystems, Foster City, CA). RNA (2 µg) was subjected to RNA sequencing using an Illumina HiSeq2000 (Otogenetics, Norcross, GA, USA). Fastq files were imported into *Partek^TM^ Flow^TM^ RNA-Seq Toolkit* software, v12.2.1. Analysis was conducted as previously described (22).

### Quantitative real time polymerase chain reaction

cDNA from TP-3654 treated cells were used for relative quantification by RT–PCR analyses. First-strand cDNA synthesis was performed from 1 μg RNA in a 20 μl reaction mixture using the high-capacity cDNA Reverse Transcription Kit (Applied Biosystems, Foster City, CA). *DDIT4*, *DDIT3*, *PPP1R15A*, *HSPA5*, *PMAIP1*, and *GAPDH* transcripts were amplified using commercially available TaqMan^®^ Gene expression assays (Applied Biosystems, Foster City, CA).

### Quantification of drug-induced cytotoxicity

Cell viability was assessed by MTT assay. Cells were seeded into 96-well microculture plates at 10,000 cells per well and allowed to attach for 24 h. Cells were then treated with TP-3654, CQ, and combinations for 72 h. Following drug treatment, MTT was added and cell viability was quantified using a Molecular Devices microplate reader. Pro-apoptotic effects following *in vitro* drug exposure were quantified by propidium iodide (PI) staining and fluorescence-activated cell sorting (FACS) analysis of sub-G_0_/G_1_ DNA content as previously described (23) and by measurement of active caspase-3 by flow cytometry using a commercial kit (BD Biosciences, Franklin Lakes, NJ).

### Measurement of ATP

Equal number of cells were plated and treated with varying concentrations of TP-3654 for 72 h. ATP levels were determined using the ATPlite assay kit, which measures the reaction of ATP with luciferase and D-luciferin. The assay was performed by following the manufacturer’s guidelines (Perkin Elmer, Waltham, MA) as previously described (24–26).

### Immunoblotting

Renal cancer cells were treated with varying concentrations of TP-3654 for 24 h. Cells were harvested and were then lysed as previously described (27). Approximately 50 μg of total cellular protein from each sample were subjected to SDS-PAGE, proteins were transferred to nitrocellulose membranes, and the membranes were blocked with 5% nonfat milk in a Tris-buffered saline solution containing 0.1% Tween-20 for 1 h. The blots were then probed overnight at 4 °C with primary antibodies, washed, and probed with species-specific secondary antibodies coupled to HRP. Immunoreactive material was detected by using the ProteinSimple ChemiGlow Chemiluminescence kit (ProteinSimple, San Jose, CA).

### Transmission electron microscopy

Transmission electron microscopy of RCC cells was performed as previously described (28). Briefly, control RCC cells or cells treated with TP-3654 or with silenced PIM1 were harvested for imaging. Sections were cut in an LKB Ultracut microtome (Leica), stained with uranyl acetate and lead citrate, and examined in a JEM 1230 transmission electron microscope (JEOL, USA, Inc.). Images were captured using the AMT Imaging System (Advanced Microscopy Techniques Corp). Manual counting was used to calculate the number of autophagosomes per cell.

### Immunocytochemistry

Cells were plated on chamber slides and allowed to attach overnight. Cells were then treated for 24 h with TP-3654. Following drug treatment, cells were fixed with 4% paraformaldehyde, permeabilized using 0.2% triton-X-100, and incubated overnight with anti-LC3B antibody. Alexa Fluor 594 or Alexa Fluor 488 conjugated fluorescent secondary antibody was used to visualize protein localization. DAPI was utilized to stain the nucleus. Images were captured using an Olympus fluorescent microscope with a DP71 camera and a 40X objective. Image-Pro Plus software Version 6.2.1 (MediaCybernetics, Bethesda, MD) was used for image acquisition (29).

### shRNA silencing of *PIM1*, *ATG7*, and *DDIT4*

In accordance with the manufacturer’s guidelines (Santa Cruz Biotechnology, Santa Cruz, CA), 786-O cells were infected with lentiviral particles containing non-targeted (control) or target-specific short hairpin RNA (shRNA) directed at *PIM1*, *ATG7*, and *DDIT4.* Effective transfection was selected for with puromycin. Transfected cells were treated with the designated drugs and concentrations. Immunoblotting was used to determine knockdown efficiency.

### Overexpression of *DDIT4*

786-O cells were infected with lentiviral particles containing GFP and a puromycin resistance gene or a *DDIT4* cDNA construct according to the manufacturer’s protocol (Origene, Rockville, MD). Successful transfection was selected for with puromycin. Cell viability and active caspase-3 were assessed as described above. Overexpression efficiency was confirmed by immunoblotting.

### Synergy Analysis

The combination indices (CI) for TP-3654 and CQ were calculated using 72 hour MTT assays. CompuSyn software (Combosyn, Inc., Paramus, NJ) was utilized to calculate CI values. In addition, synergy scores were determined using the Highest Single Agency (HSA) method as previously described (30).

### *In vivo* evaluation of TP-3654 and CQ

The animal experiments were performed with the approval of the Institutional Animal Care and Use Committee of the University of Arizona (16-094) and were conducted in accordance with established animal welfare guidelines. 786-O renal cancer cells (5 x 10^6^) were suspended in a mixture of HBSS and Matrigel and subcutaneously implanted into female nude mice (BALB/c background). Tumor-bearing animals from each cell line xenograft were randomized into treatment groups. Mice were treated with vehicle, TP-3654 (200 mg/kg PO), CQ (60 mg/kg IP), or the combination QDx5 for 6 weeks. Mice were monitored daily, and tumor volumes were measured twice weekly. At study completion, tumors from representative animals were excised from each group, formalin-fixed, and paraffin-embedded for immunohistochemical analysis.

### Immunohistochemistry

Paraffin-embedded tumor sections were deparaffinized in xylene, exposed to a graded series of alcohol, and rehydrated in PBS (pH 7.5). Heat-induced epitope retrieval on paraffin-embedded sections and probing with specific antibodies was conducted as previously described (31). Positive reactions were visualized using 3,3’-diaminobenzidine (Dako). Images were captured using an Olympus fluorescent microscope with a DP71 camera and a 20X objective. Image-Pro Plus software Version 6.2.1 (MediaCybernetics, Bethesda, MD) was used for image acquisition. ImageJ software was used for quantification of REDD1 and p62 levels by densitometric analysis of five random fields containing viable tumor cells as previously described (31). Quantification of cleaved caspase-3 and PCNA was conducted by counting the number of positive cells in five random fields as previously described (32).

### Statistical analyses

Statistical significance of differences among samples were determined using the Student’s *t* test and one-way ANOVA analysis where appropriate. Differences were considered significant in all experiments at p < 0.05.

## Results

### Inhibition of PIM kinase activity with TP-3654 selectively reduces RCC cell viability

We first treated a panel of 6 RCC cell lines (Achn, A498, 786-0, RCC4, Caki-1 and Caki-2) and normal renal proximal tubule epithelial cells (RPTEC) with various concentrations of the clinical PIM kinase inhibitor TP-3654 (chemical structure shown in **Suppl Fig. 1A**) for 72 hours. The *in vitro* effects of drug treatment on cell viability were quantified by MTT assay **(Fig. 1A)**. TP-3654 exhibited similar dose-dependent effects against the RCC cell line panel. In contrast, normal RPTEC cells were significantly less sensitive to TP-3654 than RCC cells indicating favorable therapeutic selectivity. ATPLite-mediated quantification of cellular ATP levels confirmed the dose-dependent activity of TP-3654 against RCC cells **(Fig. 1B)**. Consistent with prior studies characterizing the effects of PIM kinase inhibition, immunoblotting analyses demonstrated that TP-3654 treatment led to a dose-dependent increase in p27 expression and a decrease in the PIM kinase targets phospho-BAD and phospho-c-MYC in 786-0, A498, Achn and Caki-2 RCC cell lines **(Suppl Fig. 1B)**. Inhibition of PIM kinase activity with TP-3654 also resulted in a dose-dependent induction of apoptosis as evidenced by quantification of active caspase-3 positive cells and DNA fragmentation by PI/FACS analyses **(Suppl Fig. 2A-B)**.

**Figure 1.**
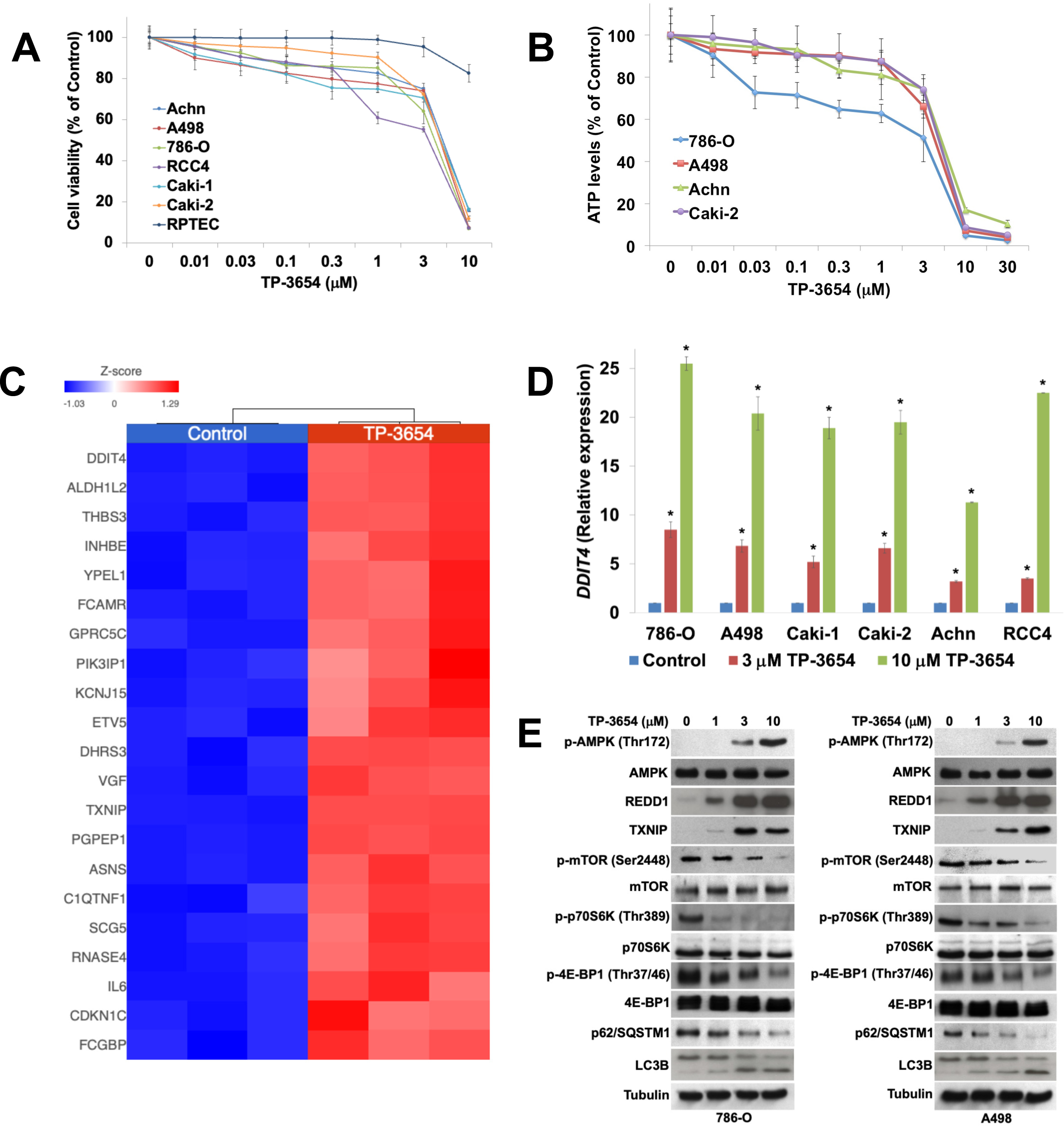
TP-3654 has selective anti-RCC effects and potently induces REDD1 expression. (**A**) TP-3654 selectively antagonizes RCC cell viability. Normal RPTEC renal cells and a panel of 6 RCC cell lines (Achn, A498, 786-O, RCC4, Caki-1 and Caki-2) were treated with the indicated concentrations of TP-3654 for 72 h. Cell viability was determined by MTT assay. Mean ± SD, n = 3. (**B**) TP-3654 diminishes ATP levels in RCC cells. 786-O, A498, Achn, and Caki-2 cells were treated with the indicated concentrations of TP-3654 for 72 h. Cellular ATP levels were quantified using the ATPLite assay according to the manufacturer’s directions. Mean ± SD, n = 3. (**C**) Effects of TP-3654 treatment on the transcriptome of RCC cells. 786-O cells were treated with 3 μM TP-3654 for 24 h. Cells were subjected to RNASeq analyses. Heatmap depicts the most significantly induced genes following TP-3654 treatment. *DDIT4* (REDD1) emerged as a highly upregulated gene in response to treatment with TP-3654. (**D**) Validation of the induction of *DDIT4* by TP-3654. 786-O, A498, Caki-1, Caki-2, Achn, and RCC4 cells were treated with the indicated concentrations of TP-3654 for 24 h. Gene expression levels were quantified using qRT-PCR and normalized to GAPDH. Mean ± SD, n = 3, * indicates significant difference from Control, p < 0.05. (**E**) Pharmacodynamic effects of TP-3654 on the AMPK/mTOR signaling cascade. 786-O and A498 cells were treated with the indicated concentrations of TP-3654 for 24 h. Protein lysates were subjected to immunoblotting to assess the effects of drug treatment on the expression of the following factors: phospho-AMPK (Thr172), total AMPK, REDD1, TXNIP, phospho-mTOR (Ser2448), total mTOR, phospho-p70S6K (Thr389), total p70S6K, phospho-4E-BP1 (Thr37/46), total 4E-BP1, p62/SQSTM1, and LC3B. Tubulin documented equal protein loading.

### Transcriptome analyses identify REDD1 (Regulated in Development and DNA Damage Responses, *DDIT4*) as a key pharmacodynamic target induced by TP-3654 treatment

To further investigate the pharmacodynamic effects of TP-3654 in RCC cells, we conducted transcriptome analyses. 786-0 cells were treated with 3 µM TP-3654 for 24 h and subjected to RNASeq. Quantification of the pharmacodynamic changes on global gene expression stimulated by TP-3654 revealed that *DDIT4* (REDD1) was one of the most highly induced genes following drug treatment (5.59 fold increase, p = 1.01E-137), **Fig. 1C, Suppl Tables 1-2)**. qRT-PCR assays confirmed that treatment with TP-3654 for 24 h yielded potent and dose-dependent induction of *DDIT4* in 6 RCC cell lines (**Fig. 1D)**. Validation of TP-3654 induced upregulation of additional genes related to endoplasmic reticular (ER) stress (*DDIT3*, *PPP1R15A*, *HSPA5*, and *PMAIP1*) using qRT-PCR is presented in **Suppl Fig. 3**.

### TP-3654 treatment leads to REDD1-linked AMPK activation and inhibition of mTOR signaling

REDD1 is an essential regulator of mTORC1 activity in response to hypoxia and metabolic stress and inactivates mTORC1 in a tuberous sclerosis complex (TSC)1/2-dependent manner (33). We next investigated the impact of PIM inhibition on the AMPK-REDD1 signaling axis. 786-0 and A498 RCC cells were treated with the indicated concentrations of TP-3654 for 24 h and the resulting pharmacodynamic effects on key factors were assessed by immunoblotting. Inhibition of PIM kinase activity with TP-3654 resulted in a dose-dependent increase in phospho-AMPK (Thr172) in a manner that corresponded with REDD1 and TXNIP induction and downregulation of mTOR activity as assessed by phospho-mTOR (**Fig. 1E**). The antagonistic effects of drug treatment on mTOR signaling were validated by assessing the impact of TP-3654 treatment on the downstream targets phospho-p70S6K and phospho-4E-BP1. As expected, based on the role of mTOR signaling as a key regulator of autophagic dynamics, treatment of 786-0 and A498 cells with TP-3654 also led to a dose-dependent reduction in p62/SQSTM1 levels and increased expression of LC3B, both of which are consistent with elevated autophagic degradation activity (**Fig. 1E**).

### Pharmacological or genetic inhibition of PIM kinase activity stimulates autophagy

We further explored the mechanistic relationship between PIM kinase activity and autophagy. 786-0 and A498 cells were treated with 3 µM TP-3654 for 24 h and stained with an anti-LC3B specific antibody. Fluorescent imaging was used to quantify the impact of TP-3654 treatment on the formation of LC3B punctae. This demonstrated that inhibition of PIM activity led to significantly higher numbers of LC3B punctae per cell (**Fig. 2A**), which is consistent with elevated autophagy activity. Transmission electron microscopy analyses also showed that TP-3654 treatment led to significantly increased mean number of autophagosomes per cell (**Fig. 2B**). Bafilomycin clamp experiments followed by immunoblotting of p62/SQSTM1 and LC3B demonstrated that TP-3654 stimulated autophagic flux (**Fig. 2C**).

**Figure 2.**
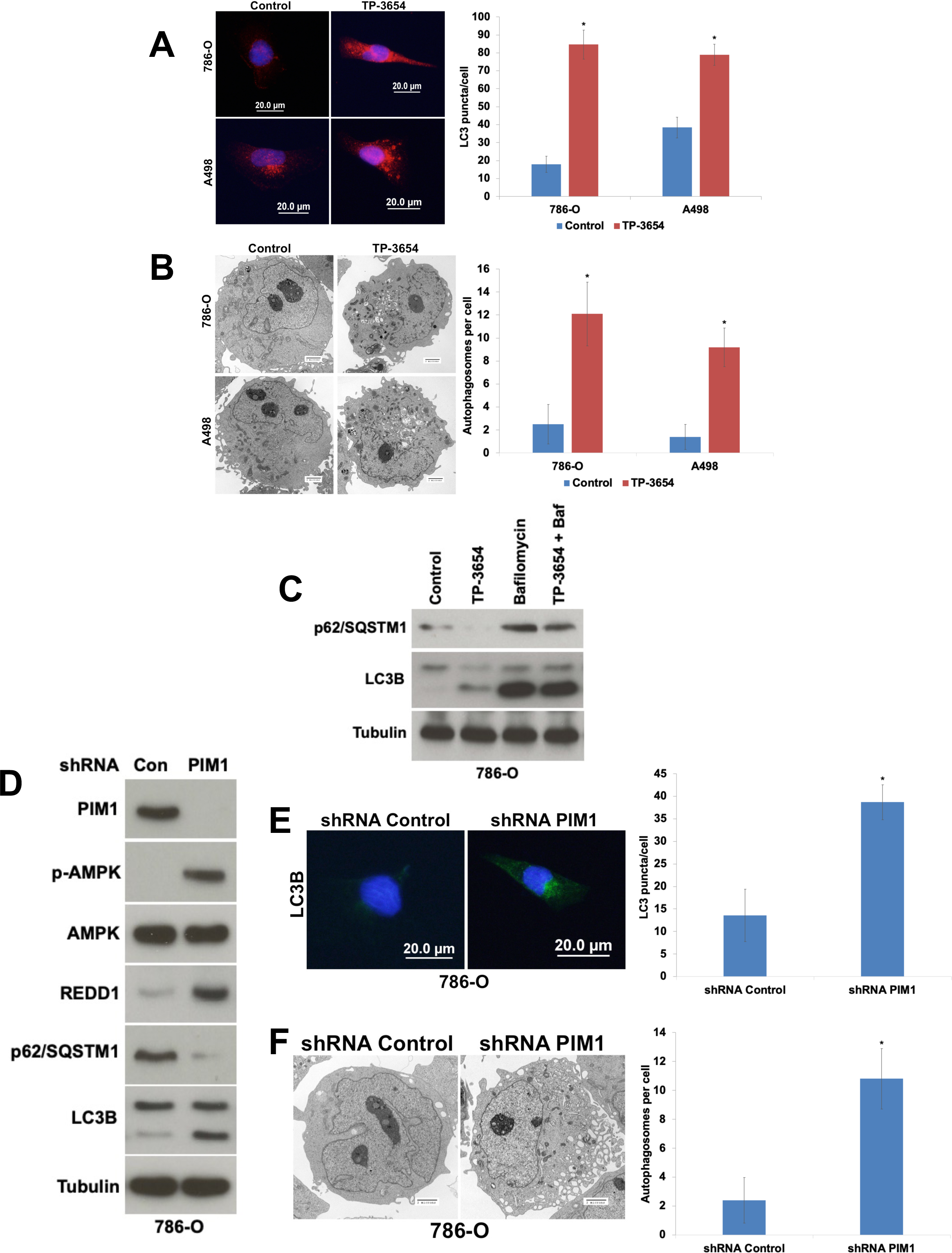
Pharmacological or genetic inhibition of PIM kinase activity stimulates autophagy. (**A**) TP-3654 induces LC3B punctae formation. 786-O and A498 RCC cells were treated with 3 μM TP-3654 for 24 h. LC3B punctae were visualized and quantified by immunocytochemistry. Mean ± SD, n = 10. (**B**) Treatment with TP-3654 triggers autophagosome formation. 786-O and A498 cells were treated 3 μM TP-3654 for 24 h. Autophagosomes were visualized and quantified using transmission electron microscopy. Mean ± SD, n = 10. (**C**) TP-3654 stimulates autophagic flux. 786-O cells were treated with 3 μM TP-3654, 100 nM bafilomycin A1, or both agents for 24 h. The effects of drug treatment on p62/SQSTM1 and LC3B protein expression were determined by immunoblotting. Tubulin documented equal protein loading. (**D**) Genetic inhibition of *PIM1* activates the AMPK-REDD1-autophagy cascade. Lentiviral shRNA was used to silence *PIM1* in 786-O cells. Knockdown efficiency was evaluated by immunoblotting. The effects of genetic *PIM1* inhibition on AMPK activation, REDD1 expression, and autophagy (p62/SQSTM1 and LC3B) were assessed by immunoblotting. (**E**) Targeting *PIM1* induces LC3B punctae formation. Basal LC3B punctae in 786-O cells infected with control or *PIM1*-targeted shRNA were visualized and quantified by immunocytochemistry. Mean ± SD, n = 10. (**F**) Genetic impairment of *PIM1* induces autophagosome formation. 786-O cells were infected with control or PIM1-targeted shRNA and autophagosomes were visualized and quantified using transmission electron microscopy. Mean ± SD, n = 10. * indicates a significant difference from controls, p < 0.05.

All drugs have off-target effects that vary based on the specific agent and the particular dose that is used. To rule out the possibility that the induction of REDD1 and autophagy that is stimulated by treatment with TP-3654 could be driven by a PIM-independent drug effect, we utilized shRNA to genetically impair *PIM1*. Targeted knockdown of PIM1 expression in 786-O cells recapitulated all of the key pharmacodynamic effects we observed with pharmacological PIM kinase inhibition including phosphorylation of AMPK, upregulation of REDD1, and activation of autophagy (decreased p62 levels and increased LC3B expression) as shown by immunoblotting analyses (**Fig. 2D**). Accordingly, *PIM1* shRNA also significantly increased the number of LC3B punctae per cell (**Fig. 2E**) and the mean number of autophagosomes per cell (**Fig. 2F**). Collectively, these data establish a link between PIM inhibition, REDD1 upregulation, and the stimulation of autophagy.

Autophagy is a well-established mechanism of resistance to multiple classes of anticancer agents (34,35). To further investigate whether the pharmacodynamic autophagy activation associated with PIM inhibition significantly impacts therapeutic sensitivity, we utilized shRNA targeting the essential autophagy gene *ATG7* to genetically impair the pathway in 786-O cells (**Fig. 3A**). Treatment of control and *ATG7* shRNA infected 786-O cells with TP-3654 demonstrated that disrupting autophagy significantly increased the anti-RCC benefit of PIM kinase inhibition with respect to diminishing cell viability and inducing apoptosis (**Fig. 3B**). If there is a robust relationship between PIM function and autophagic activity, then we hypothesized that impairing PIM would be expected to increase sensitivity to autophagy inhibition. We tested this mechanistic relationship using *PIM1*-targeted shRNA in 786-O cells (**Fig. 3C**). Indeed, targeted PIM1 knockdown significantly increased the *in vitro* anticancer activity of chloroquine (CQ) in a dose-dependent manner (**Fig. 3D**).

**Figure 3.**
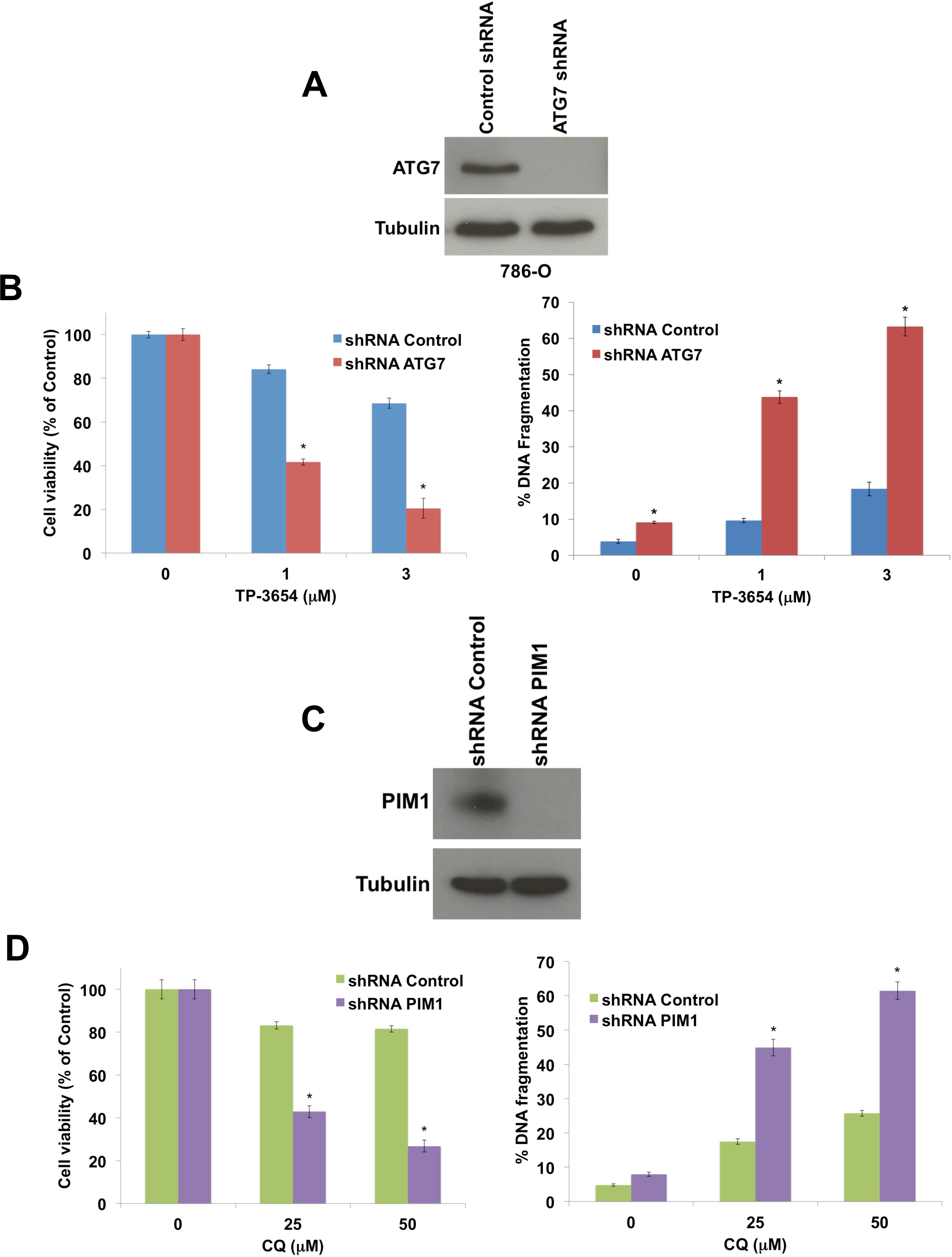
Genetic *PIM1* and autophagy inhibition cooperate to enhance anti-RCC activity. (**A**) The essential autophagy gene *ATG7* was knocked down in 786-O RCC cells using lentiviral shRNA. Knockdown efficiency was assessed by immunoblotting. Tubulin documented equal protein loading. (**B**) Cells infected with non-targeted control or *ATG7*-directed lentiviral shRNAs were treated with the indicated concentrations of TP-3654 for 72 h (left, MTT) or 48 h (right, PI/FACS). The effects of drug treatment on cell viability (left) and apoptosis (right) were quantified for each experimental condition. Mean ± SD, n = 3. (**C**) *PIM1* was knocked down in 786-O RCC cells using lentiviral shRNA. Knockdown efficiency was assessed by immunoblotting. (**D**) Targeted *PIM1* knockdown enhances the anti-RCC activity of CQ. Cells infected with non-targeted control or *PIM1*-directed lentiviral shRNAs were treated with the indicated concentrations of CQ for 72 h (left, MTT) or 48 h (right, PI/FACS). The effects of drug treatment on cell viability (left) and apoptosis (right) were quantified for each experimental condition. Mean ± SD, n = 3. * indicates a significant difference from controls, p < 0.05.

### REDD1 expression is a key determinant of sensitivity to autophagy inhibition

Our initial studies exploring the mechanism of action of TP-3654 in RCC cells showed that PIM kinase inhibition leads to a dramatic upregulation of REDD1 that is linked to autophagy activation and increased sensitivity to autophagy inhibition. To specifically investigate the role of REDD1 (*DDIT4*) as a regulator of sensitivity to autophagy inhibitors, we utilized shRNA to knockdown *DDIT4* in 786-O RCC cells in addition to isogenic HAP1 cells with and without *DDIT4* knockout (**Fig. 4A**). Genetic impairment of *DDIT4* expression dramatically blunted the ability of CQ to induce LC3 punctae (**Fig. 3B**) and reduced cellular sensitivity to CQ-mediated autophagy inhibition (**Fig. 3C-D**). Conversely, overexpression of REDD1 significantly augmented the ability of CQ to reduce RCC cell viability and induce apoptotic cell death (**Fig. 4E** and **Supp. Fig. 4**). Collectively, these findings demonstrate that REDD1 expression is an important determinant of the sensitivity of RCC cells to autophagy inhibition.

**Figure 4.**
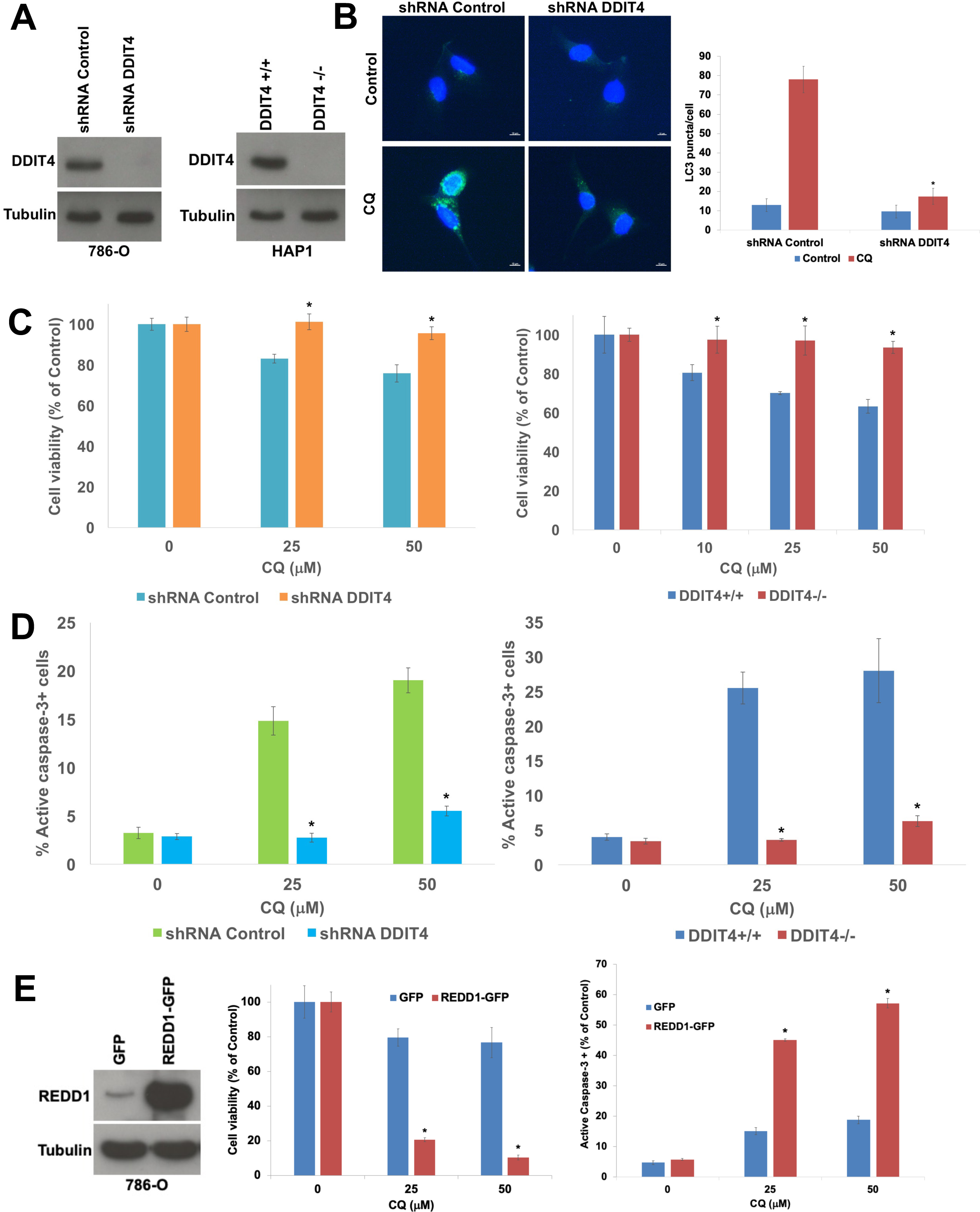
*DDIT4*/REDD1 status determines sensitivity to autophagy inhibition. (**A**) *DDIT4* was knocked down in 786-O RCC cells using lentiviral shRNA and knocked out in HAP1 cells using CRISPR. Knockdown/knockout efficiency was assessed for both cell lines by immunoblotting. (**B**) *DDIT4* knockdown significantly reduces CQ-induced LC3B punctae formation. 786-O cells infected with control or *DDIT4*-targeted shRNA were treated with 25 µM CQ for 24 h. LC3B punctae were visualized and quantified by immunocytochemistry. Mean ± SD, n = 10. * indicates a significant difference from shRNA controls, p < 0.05. (**C**) Genetic impairment of *DDIT4* reduces cellular sensitivity to CQ. 786-O cells infected with control or *DDIT4*-targeted shRNA (left) and isogenic HAP1 cells with and without *DDIT4* knockout (right) were treated with the indicated concentrations of CQ for 72 h. The impact of drug treatment on cell viability was determined by MTT assay. Mean ± SD, n = 3. * indicates a significant difference from shRNA controls or *DDIT4* +/+ cells, p < 0.05. (**D**) Targeting *DDIT4* blunts the pro-apoptotic effects of CQ. 786-O cells infected with control or *DDIT4*-targeted shRNA (left) and isogenic HAP1 cells with and without *DDIT4* knockout (right) were treated with the indicated concentrations of CQ for 48 h. The impact of drug treatment on apoptosis induction was determined by staining with a FITC-tagged active caspase-3 antibody followed by flow cytometry. Mean ± SD, n = 3. * indicates a significant difference from shRNA controls or *DDIT4* +/+ cells, p < 0.05. (**E**) REDD1 (*DDIT4*) overexpression synergistically enhances the sensitivity of RCC cells to autophagy inhibition. REDD1 was overexpressed in 786-O cells using a GFP-tagged lentiviral construct. Overexpression efficiency was assessed by immunoblotting (left). GFP and REDD1-GFP overexpressing cells were treated with the indicated concentrations of CQ for 72 h (middle, MTT) or 48 h (right, active caspse-3). The effects of drug treatment on cell viability (middle) and apoptosis (right) were quantified for each experimental condition. Mean ± SD, n = 3. * indicates a significant difference from GFP control cells, p < 0.05.

### Co-targeting PIM kinase activity and autophagy yields synergistic anti-RCC effects

We next explored the potential benefit of co-targeting PIM kinase activity and autophagy for RCC therapy. Treatment of 786-O, A498, Achn and Caki-2 cells with the combination of TP-3654 and CQ led to a dramatically greater reduction in cell viability than either single agent by MTT assay (**Fig. 5A**) and quantification of cellular ATP levels (**Fig. 5B**). Formal synergy analyses demonstrated that the combination of TP-3654 and CQ synergistically reduced cell viability in all 4 models that were evaluated (**Fig. 5C** and **Suppl Table 3**). Accordingly, co-treatment with TP-3654 and CQ also resulted in significantly greater levels of apoptosis induction than either single agent in a panel of 6 RCC cell lines (**Fig. 5D**). The synergistic pro-apoptotic effects of this combination strategy were further supported by the significantly higher levels of PARP cleavage in 786-O and A498 cells (**Fig. 5E**). Taken together, these data indicate that simultaneous inhibition of PIM kinase activity and autophagy yields synergistic benefits against RCC cells.

**Figure 5.**
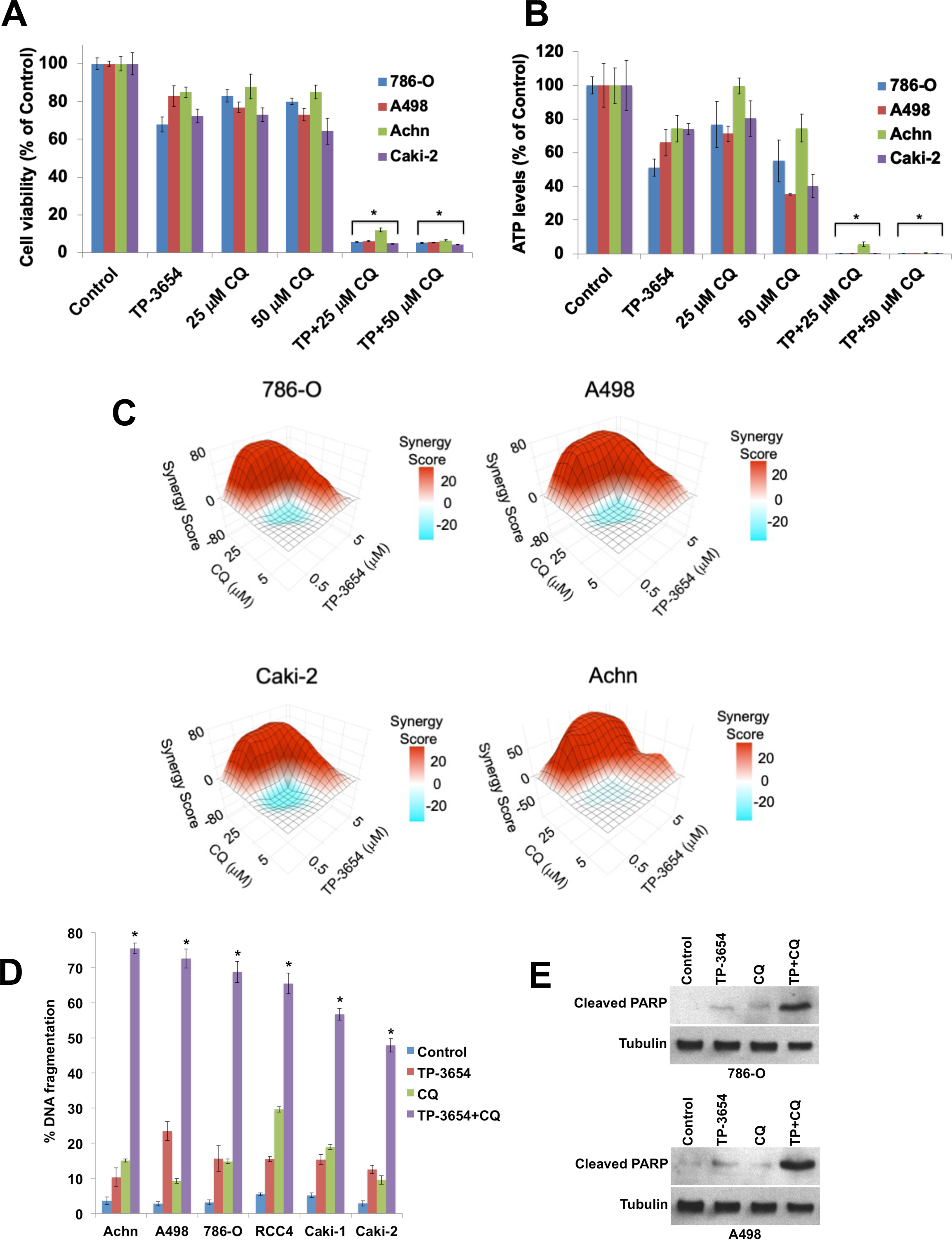
Inhibition of autophagy with CQ synergistically enhances the anti-RCC effects of TP-3654. (**A**) A panel of 4 RCC cell lines (Achn, A498, 786-O and Caki-2) were treated with 3 µM TP-3654 alone, the indicated concentrations of CQ, or both drugs for 72 h. Cell viability was determined by MTT assay. Mean ± SD, n = 3. (**B**) The TP-3654/CQ combination synergistically diminishes ATP levels in RCC cells. 786-O, A498, Achn, and Caki-2 cells were treated with 3 µM TP-3654, the indicated concentration of CQ, or both drugs for 72 h. Cellular ATP levels were quantified using the ATPLite assay according to the manufacturer’s directions. Mean ± SD, n = 3. (**C**) Heatmaps of the synergistic effects of the TP-3654/CQ combination. 786-O, A498, Caki-2 and Achn cells were treated with varying concentrations of TP-3654, CQ, or both drugs for 72 h. The effects of each treatment on cell viability were determined by MTT assay and the resulting data was utilized to establish a synergy score for the combination in each cell line. The heatmaps depict the synergy dynamics across all drug concentrations that were evaluated. (**D**) TP-3654 and CQ have broad and synergistic pro-apoptotic effects against RCC cells. A panel of 6 RCC cell lines (Achn, A498, 786-O, RCC4, Caki-1 and Caki-2) were treated with 3 µM TP-3654, 25 µM CQ, or the combination for 48 h. PI/FACS analysis was used to quantify the percentages of cells with fragmented DNA as a measure of apoptosis for each treatment condition. Mean ± SD, n = 3. * indicates a significant difference from controls and either single agent treatment, p < 0.05. (**E**) The TP-3654/CQ combination synergistically stimulates PARP cleavage. 786-O and A498 cells were treated with 3 µM TP-3654, 25 µM CQ, or the combination for 24 h. Immunoblotting was used to determine the effects of drug treatment on PARP cleavage. Tubulin documented equal protein loading.

### Administration of the TP-3654/CQ combination to mice bearing RCC xenografts is well tolerated and yields synergistic tumor regression

Finally, we conducted a mouse xenograft study to evaluate the synergistic efficacy of the TP-3654/CQ combination *in vivo*. Immunodeficient nude mice bearing 786-O RCC tumors were randomized into vehicle, TP-3654, CQ, or the TP-3654/CQ combination treatment groups. Drug treatment continued for 6 weeks and disease burden was monitored twice weekly. TP-3654 and CQ monotherapies had modest effects curtailing tumor progression. In contrast, simultaneous inhibition of PIM kinase activity and autophagy with the TP-3654/CQ combination led to stable tumor regression in a manner that mirrored the synergy we observed in our *in vitro* studies with these agents (**Fig. 6A**). All treatments were very well tolerated over the course of the study and yielded a negligible impact on mouse weight (**Fig. 6B**). Pharmacodynamic analyses of tumor specimens collected from mice in each treatment group demonstrated that inhibition of PIM kinase activity with TP-3654 potently induced REDD1 levels *in vivo* (**Fig. 6C**). Administration of CQ effectively blocked the degradation of p62 that was driven by TP-3654-induced autophagy. Quantification of the effects of drug administration on PCNA and active caspase-3 levels in 786-O tumors showed that the TP-3654/CQ combination more effectively inhibited tumor cell proliferation (PCNA) and induced apoptosis (active caspase-3) than either single agent therapy (**Fig. 6C**). Our collective data identify REDD1 as a key regulator of the sensitivity of RCC cells to autophagy inhibition. We demonstrate a mechanistic link between PIM kinase activity, REDD1 levels, and autophagic degradation. Our findings establish co-targeting PIM kinases and autophagy as a well-tolerated approach to therapeutically exploit the PIM-REDD1-autophagy axis that warrants further investigation (**Suppl Fig. 5**).

**Figure 6.**
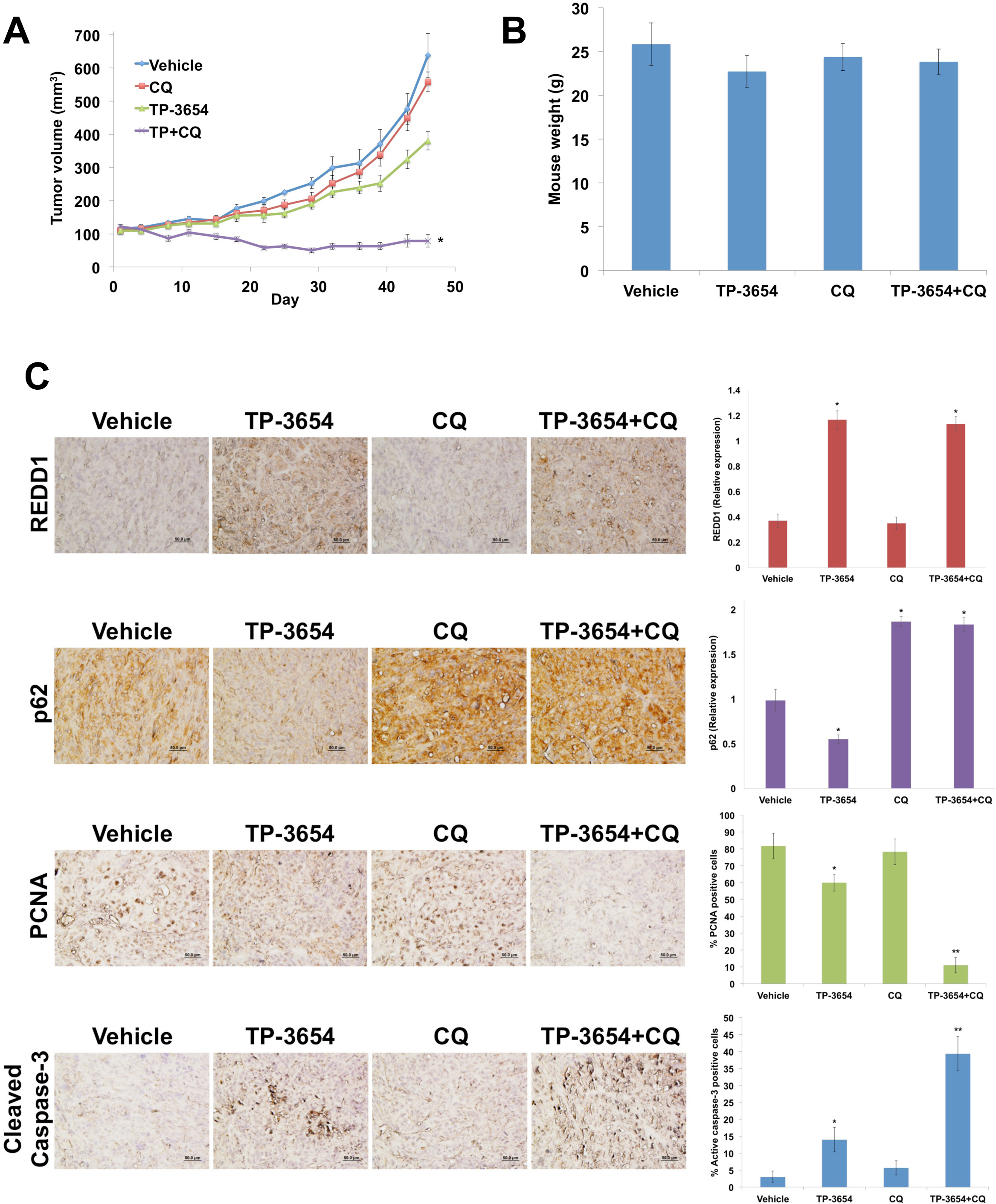
Co-targeting PIM kinase activity and autophagy is well tolerated and yields tumor regression in a RCC xenograft model. (**A**) 786-O cells were injected into the flanks of nude mice. Mice were pair-matched and randomized into groups when mean tumor burden reached approximately 100 mm^3^. Mice were treated with 200 mg/kg TP-3654 PO, 60 mg/kg CQ IP, and the combination QDx5 throughout the course of the study. Tumor volumes were measured twice weekly. Mean ± SEM, n = 6. *Indicates a significant difference between combination treated mice vs. vehicle or single agents, p < 0.05. (**B**) The TP-3654/CQ combination is well tolerated in mice. Body weight was determined at the end of the study (Day 46) to quantify drug-induced weight loss. Mean ± SD, n = 6. (**C**) Immunohistochemistry-based quantification of the pharmacodynamic effects of drug treatment. Tumors from mice in each treatment group were collected at study endpoint. Tumor sections were stained with antibodies against REDD1, p62, PCNA and cleaved caspase-3. Representative images were captured under 20X magnification. The relative intensity of expression of REDD1 and p62 was quantified by densitometry. Mean ± SD, n = 5. The percentage of cleaved caspase-3 and PCNA positive cells was determined manually under 20X magnification. Mean ± SD, n = 5.

## Discussion

Survival outcomes for patients with ccRCC have significantly improved over the last decade due to the availability of new treatment options. However, patients with advanced metastatic disease and those that develop drug resistance after receiving standard therapies continue to have an extremely poor prognosis and are in urgent need of additional treatment options (36). Targeting PIM kinase activity is a strategy that has been explored for the treatment of multiple cancer types, although it has not been previously rigorously investigated specifically for ccRCC (37). The concept of therapeutic PIM kinase inhibition is attractive as the members of this kinase family are overexpressed in a number of different malignancies and earlier genetic studies that ablated its function demonstrated strong therapeutic potential in the preclinical setting. Accordingly, multiple pharmaceutical companies including Novartis, AstraZeneca, and Sumitomo have launched programs focused on developing clinical stage PIM kinase inhibitors for cancer therapy.

Multiple PIM kinase inhibitors have advanced into clinical trials, but none have earned FDA approval to date. The reasons for this are likely multifold, but one that is undoubtedly a significant contributor to this current fate is the incomplete understanding of how PIM inhibition reprograms the signaling dynamics of malignant cells that survive this attack. We aimed to decipher this in ccRCC cells in the current investigation to empower the selection of a rational approach to improve the therapeutic benefit of PIM kinase inhibition. Our study revealed that PIM inhibition leads to the potent induction of autophagy in a manner that negatively impacts the anticancer activity of TP-3654. This is not an entirely surprising result given the connection between PIM kinases and the regulation of the AMPK-mTOR signaling cascade (38,39). However, our discovery that REDD1 plays a pivotal role in this process is novel and provided an interesting opportunity to further explore its links to the control of both PIM and autophagic activity.

We and others have previously shown that disrupting autophagy has significant promise as an approach for ccRCC therapy (5,40). In fact, a patient with ccRCC who enrolled on our phase I investigator-initiated clinical trial of HCQ in combination with vorinostat experienced an amazing long-term response lasting more than 10 years despite failing 7 prior lines of conventional and investigational therapy (31). However, the mechanistic basis for his fantastic outcome remains a mystery. Indeed, one of the greatest challenges that investigators pursuing therapeutic autophagy inhibition as an anticancer strategy have faced regardless of tumor type is the lack of a clear and validated biomarker that predicts which patients are most dependent on the pathway and therefore likely to yield greater benefit from this approach. With HCQ/CQ still being evaluated in cancer clinical trials along with second generation autophagy inhibitors ezurpimtrostat (GNS561) (41) and the ULK1/2 inhibitor DCC-3116 (42), identifying potential predictive biomarkers of sensitivity is essential. For example, previous preclinical studies have suggested that tumors with activated RAS status are specifically dependent upon autophagy (43,44). Yet, there are many patients who were treated with HCQ-based regimens in early phase clinical trials who exhibited activating RAS mutations and failed to achieve clinical responses. It is obvious that other factors are important in determining the degree of benefit that patients experience when treated with autophagy inhibitor-based regimens.

REDD1 forms a complex with TXNIP, which is sufficient to induce reactive oxygen species, suppress ATG4B activity, and stimulate autophagy (45). Consistent with this observation, we found that REDD1 deficient cells display defects in autophagy and thus, are resistant to cell death induced by autophagy inhibitors. It has been previously reported that REDD1 expression may be upregulated by HIFs under hypoxic conditions and this may explain its relatively abundant expression in ccRCC (46). However, REDD1 can also be induced by several other transcription factors including p53 and ATF4 and it is possible that these mechanisms may also contribute to the regulation of its expression in ccRCC (47–49). Here we report for the first time that REDD1 status is a determinant of the sensitivity of ccRCC cells to autophagy inhibition. This finding is significant for two reasons. First, our data establish the rationale to further investigate the relationship between REDD1 expression levels and sensitivity to autophagy inhibitors alone and in combination with other agents to determine its suitability as a candidate biomarker that predicts response to autophagy inhibition. Not only can this be examined in a prospective manner, but it could also be interrogated retrospectively through the evaluation of datasets from completed HCQ involved cancer clinical trials. Second, our study revealed that PIM kinase inhibition is a highly effective mechanism to upregulate REDD1 expression. This approach synergized with CQ to yield tumor regression in our ccRCC xenograft model, which is obviously significant from a combination therapy perspective. Beyond that, this finding offers a potential opportunity to transform “autophagy inhibitor insensitive” tumors to “autophagy inhibitor hypersensitive tumors” through targeting PIM kinase activity in a manner that is analogous to the concept of immune-reprogramming immune cold tumors to heighten their response to immunotherapy (50).

Collectively, our findings identify REDD1 as a novel determinant of the sensitivity of ccRCC cells to autophagy inhibition and establish a mechanistic link between PIM kinase activity, REDD1 status and autophagic degradation in this specific tumor type. We demonstrate that co-targeting PIM kinase activity and autophagy yields synergistic anticancer activity and tumor regression in preclinical models of ccRCC. Further investigation into the role of REDD1 as a candidate biomarker that predicts response to autophagy inhibition and of the safety and efficacy of combined PIM and autophagy inhibition is warranted.

## Financial Support

Research reported in this publication was supported by the National Cancer Institute of the National Institutes of Health under award numbers R01CA268383, T32CA009213, and P30CA023074. Support was also provided by the University of Arizona Integrative Cancer Scholars – Cancer Research Training and Education Coordination (CRTEC) fellowship.

## Conflict of Interest

JSC, WW, and STN are co-founders of Majestic Therapeutics, LLC.

## Disclosure of conflicts of interest

JSC, WW, and STN are co-founders of Majestic Therapeutics, LLC. None of the other authors have any relevant conflicts of interest to declare.

## Author contributions

Conception and design – JSC, STN

Development of methodology – JSC, CME, STN

Data acquisition, analysis, and interpretation - JSC, CME, SS, MJCE, MEG, WW, BL, STN

Manuscript preparation – JSC, WW, BL, STN

## Grant support

This work was supported by grants from the National Cancer Institute (R01CA269393, T32CA009213, P30CA023074) and the University of Arizona Cancer Center ICS-CRTEC fellowship.

