## Supplemental Figures for "REDD1 is a determinant of the sensitivity of renal cell carcinoma cells to autophagy inhibition that can be therapeutically exploited by targeting PIM kinase activity"

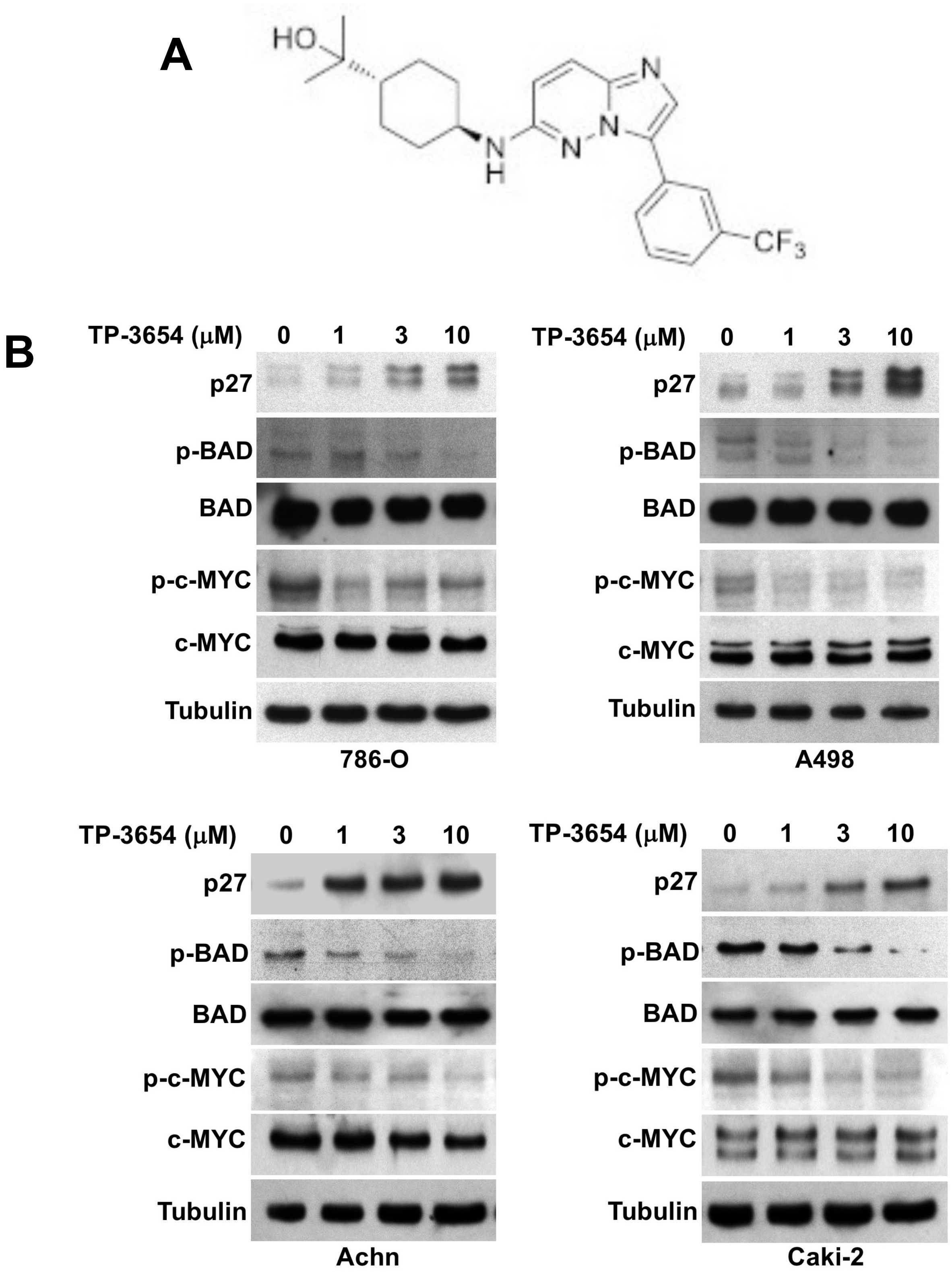


**Supplemental Figure 1.** Effects of TP-3654 on RCC cell lines. (**A**) Chemical structure of TP-3654. (**B**) TP-3654 treatment induces p27 expression and decreases the phosphorylation of BAD (Ser112) and c-MYC (Ser62). RCC cells were treated with the indicated concentrations of TP-3654 for 24 h. Protein expression was determined by immunoblotting.

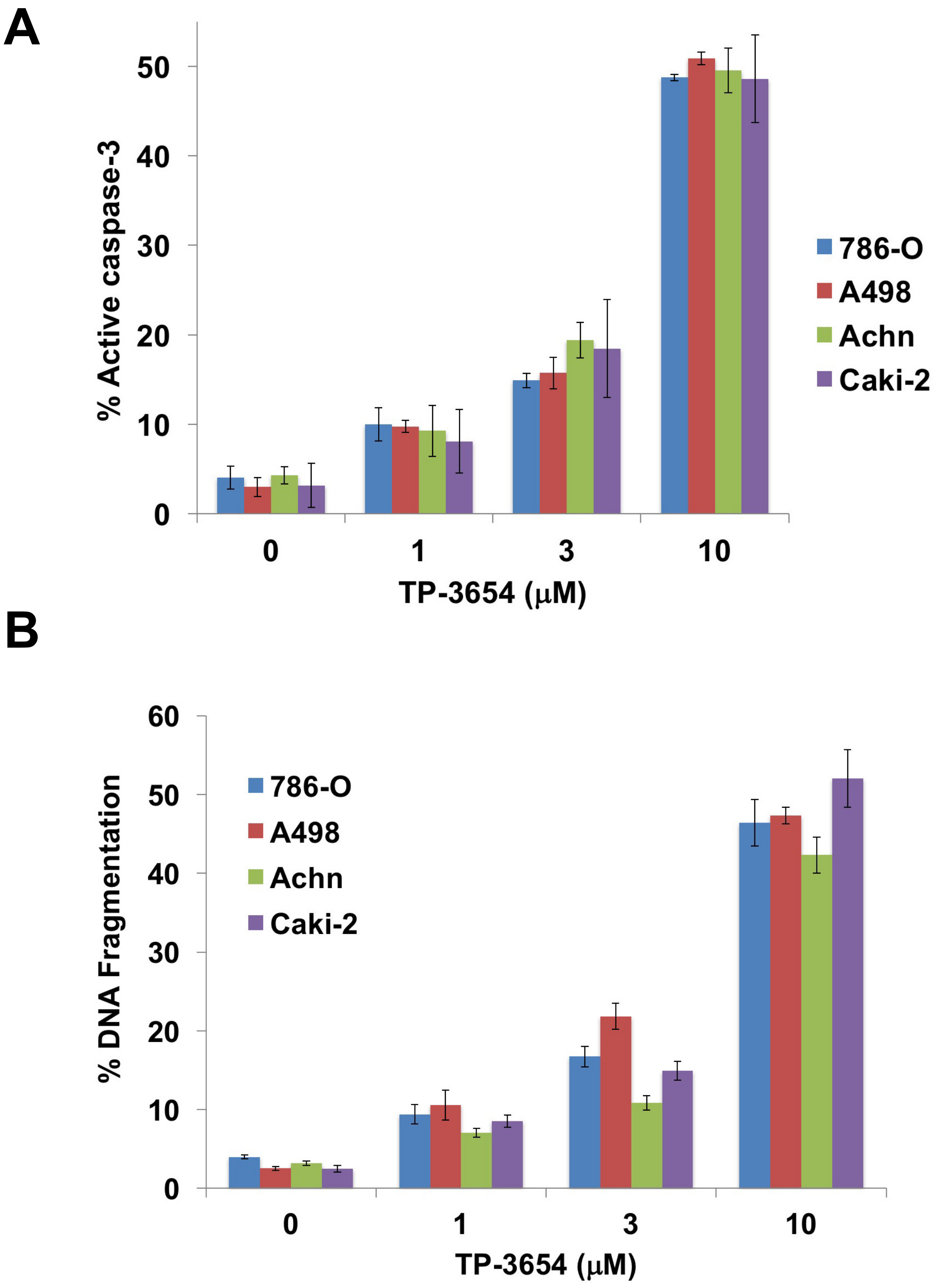


**Supplemental Figure 2.** TP-3654 induces apoptosis in RCC cells. A panel of RCC cell lines were treated with the indicated concentrations of TP-3654 for 48 h. (**A**) Active caspase-3 assay and (**B**) PI/FACS analysis were used to quantify apoptosis. Mean ± SD, n = 3.

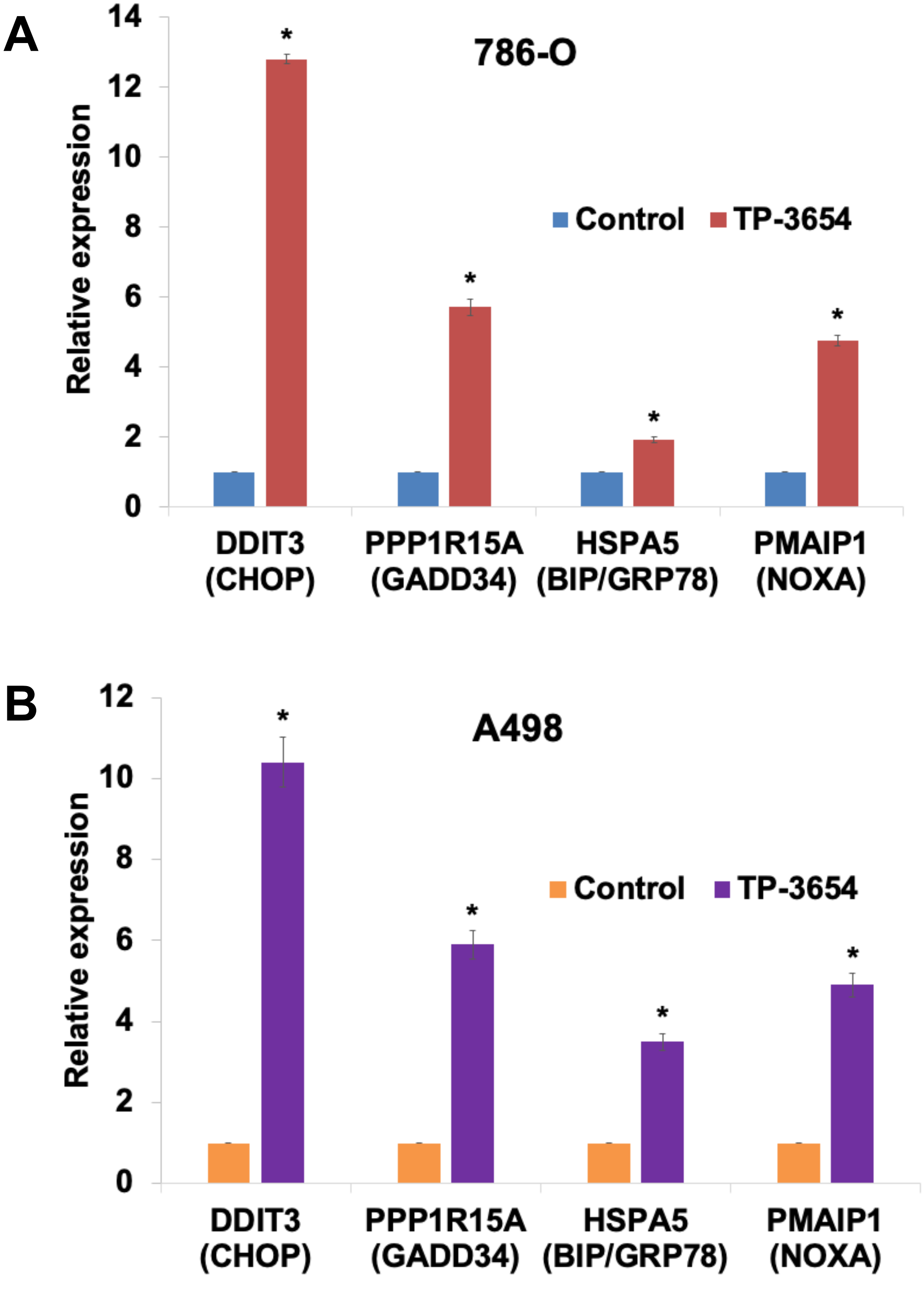


**Supplemental Figure 3.** TP-3654 induces ER stress-related gene expression. (**A-B**) 786-O and A498 cells were treated with 3 μM TP-3654 for 24 h. Gene expression levels of *DDIT3*, *PPP1R15A*, *HSPA5*, and *PMAIP1* were quantified using qRT-PCR and normalized to *GAPDH*. Mean ± SD, n = 3. * Indicates a significant difference from controls, $p < 0.05$.

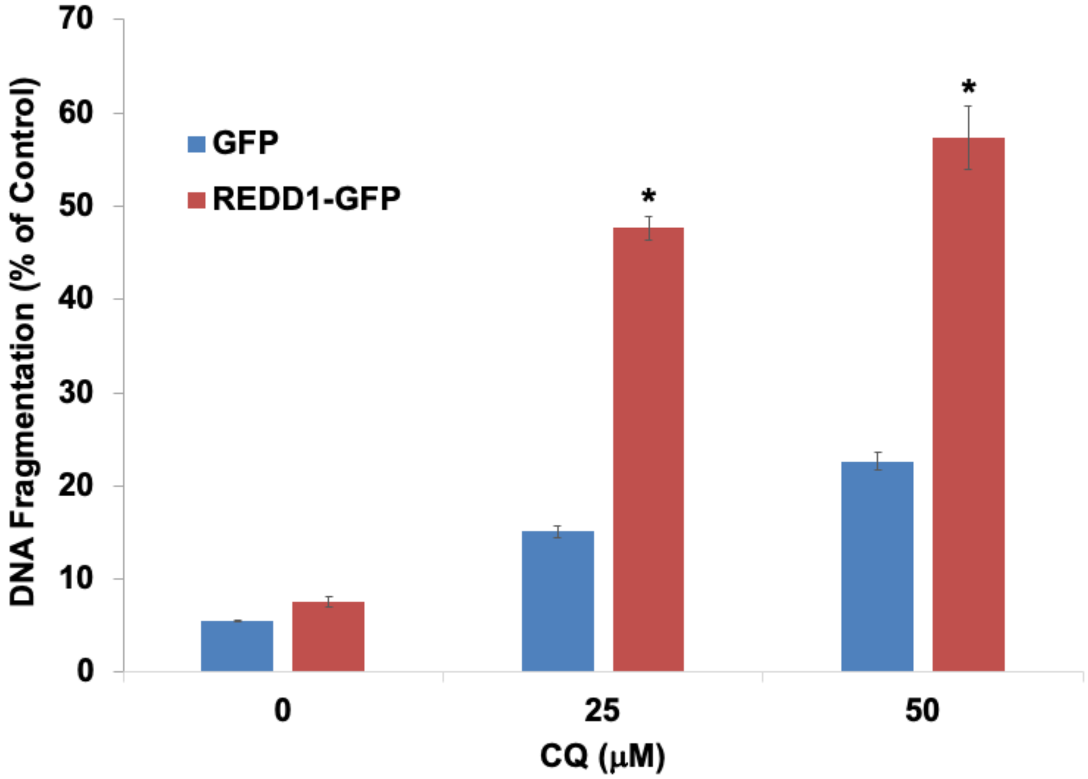


**Supplemental Figure 4.** Overexpression of REDD1 sensitizes 786-O cells to CQ-mediated apoptosis. GFP and REDD1-GFP expressing cells were treated with the indicated concentrations of CQ for 48 h. PI/FACS analysis was used to quantify the percentage of DNA fragmentation. Mean ± SD, n = 3. * Indicates a significant difference from GFP Control cells, $p < 0.05$.

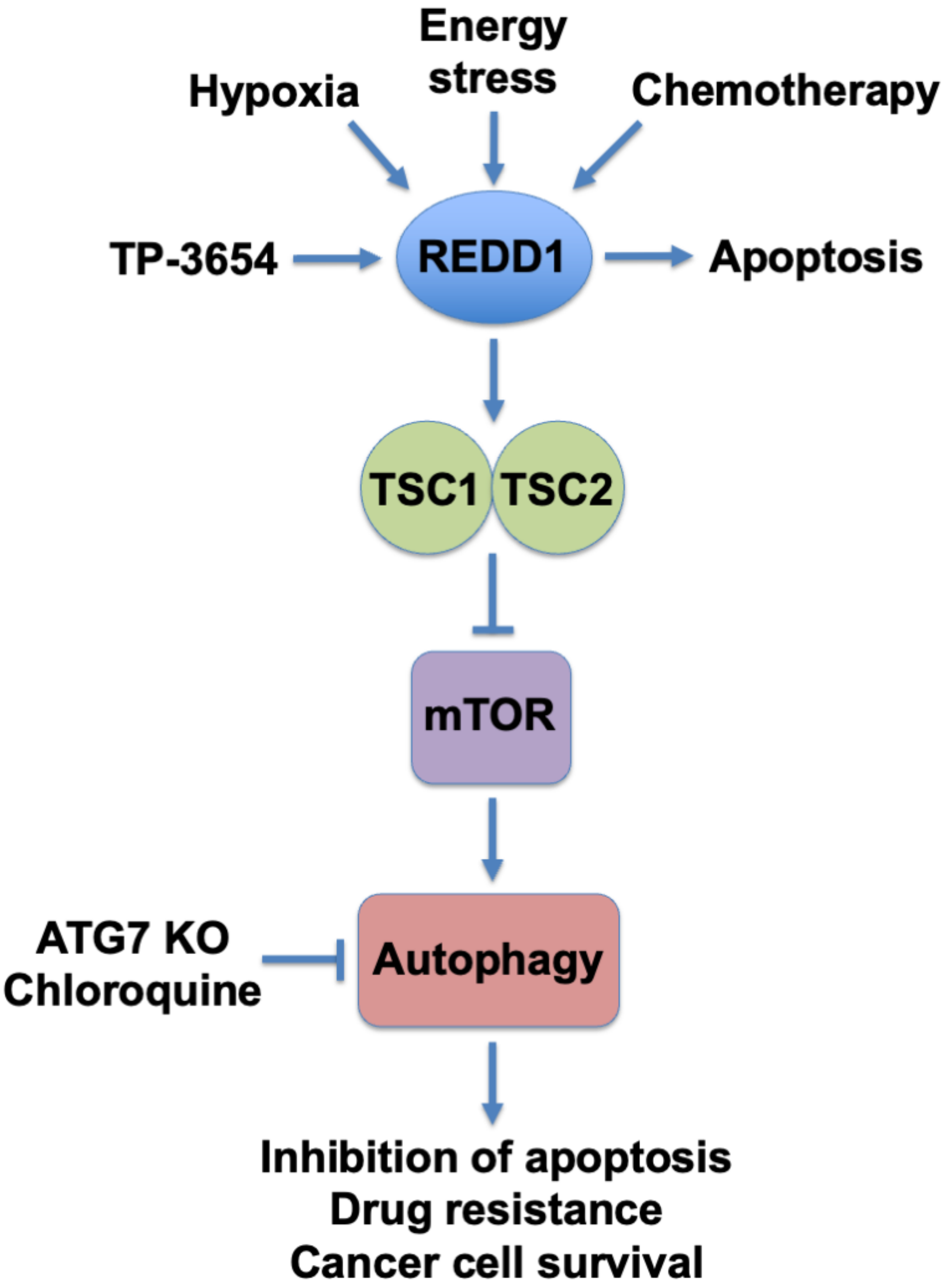


**Supplemental Figure 5.** Schematic of apoptosis induced by the combination of TP-3654 and Inhibition of autophagy. TP-3654 potently induces REDD1 expression, which results in inhibition of mTOR and upregulation of autophagy. High REDD1 expression promotes hyperactive autophagy leading to enhanced sensitivity to CQ.
